# Neuropeptide oxytocin facilitates its own brain-to-periphery uptake by regulating blood flow dynamics and permeability

**DOI:** 10.1101/2024.08.07.606980

**Authors:** Preethi Rajamannar, Oren Raz, Gil Levkowitz

## Abstract

The hypothalamo-neurohypophyseal system is an important neuroendocrine brain-to-blood conduit through which the neurohormones oxytocin and arginine-vasopressin are released from the brain into the general circulation to affect peripheral physiological functions such as salt balance, metabolism and reproduction. However, the mechanism which executes fast and efficient neurohormone release to the periphery remains unsolved. We show, using live imaging in zebrafish, that a hyperosmotic physiological challenge elicits a local increase in neurohypophyseal blood flow velocities and a change in capillary diameter, which is dictated by the geometry of the hypophyseal vascular microcircuit. Genetic ablation of oxytocin neurons and inhibition of oxytocin receptor signaling attenuated changes in capillary blood flow and diameter. Optogenetic stimulation of oxytocin neurons resulted in an oxytocin receptor-dependent increase in blood flow velocities. Lastly, both osmotic challenge and oxytocin neuronal activation elicited a local rise in neurohypophyseal capillary permeability in an oxytocin signaling-dependent manner. Our study demonstrates that physiologically elicited changes in neurohypophyseal blood flow and permeability are regulated by oxytocin. We propose that oxytocin-dependent neuro-vascular coupling facilitates its efficient uptake into the blood circulation, suggesting a self-perpetuating mechanism of peripheral hormone transfer.

## INTRODUCTION

How the brain exerts control over peripheral functions is a fundamental question that has occupied researchers for over a century. In the early 1900s, Berta and Ernest Scharrer noticed that neurons residing in the hypothalamus of a small freshwater fish have properties similar to hormone-secreting endocrine cells (*1, 2*). This discovery served as an inspiration for Scharrers’ “neurosecretion concept”, arguing that hypothalamic neurons release packets of neuro-hormones from the brain into the general circulation by directly projecting their axons onto local blood vessels of the posterior pituitary gland, also known as the neurohypophysis. This brain-to-periphery neurosecretion machinery termed the hypothalamo-neurohypophyseal system (HNS), is evolutionarily conserved among all vertebrate species and is a major neuroendocrine system responsible for the secretion of the neuro-hormones oxytocin (OXT) and arginine-vasopressin (AVP) from hypothalamic axonal termini onto fenestrated (i.e., permeable) capillary network of the neurohypophysis to affect peripheral organs. In the periphery, AVP regulates water homeostasis by increasing the water permeability of the kidney, whereas OXT regulates labor and milk let-down by causing the contraction of the uterine smooth muscles and the myoepithelial cells of breast ducts, respectively(*3, 4*). In teleost fish species, OXT and AVP are involved in osmoregulation and regulating reproductive success(*5, 6*).

Despite the physiological importance of OXT and AVP, there is a gap in our understanding of their regulated secretion into the blood. The prevailing dogma maintains that in response to physiological demands, hypothalamic OXT and/or AVP neurons are stimulated, released from the axonal termini into the neurohypophysis, taken up by blood capillaries into the systemic circulation, and transported to target tissues (*7*). This process requires tight regulation between the secretory neurons, the release of the hormones, and their transport into the blood. However, it has been noted that a simplified stimulus–secretion coupling mechanism, wherein the release of the contents of vesicles upon neuronal activation (e.g. neurotransmitter release), is directly associated with its effect on the target neuron does not apply in the context of neuroendocrine effect on peripheral targets (*8*). Thus, while the measured firing rhythms of OXT and/or AVP neurons range within milliseconds or even seconds, it takes several minutes to detect them in the peripheral blood (*9*). Moreover, unlike the dense synaptic connection between neurons or the tightly arranged neurovascular units observed in the brain, HNS nerve endings do not form compact, juxtaposed connections with pituitary blood vessels. They rather form numerous axonal varicosities packed with large dense core vesicles that release their content into the neurohypophysis, wherein the secreted neurohormones are required to transverse the cell-abundant pituitary parenchyma, the endothelial basement membrane, and then enter the endothelial lumen (*10*). ***It is, therefore, highly plausible that an additional driving force is required to facilitate efficient coupling between neurohormone release and its vascular uptake, leading to downstream effects on peripheral target tissues*.**

In the brain, synaptic activity increases local blood flow to meet the metabolic needs of a firing neuron, a process termed neurovascular coupling. It has been suggested that endocrine cells of the pituitary gland and pancreas are coordinated with blood vessels to generate hormone pulses (*11, 12*). Thus, local blood flow in neuroendocrine interfaces may facilitate efficient hormone uptake rather than just a means to cope with the increased demands for oxygen.

Here, we hypothesized that in the neurohypophysis, activity-dependent changes in local blood flow modulate peripheral neurohormone uptake. Using live imaging of the zebrafish HNS, we monitored hypophyseal vascular dynamics in unanesthetized animals. We demonstrate that the physiological activation of oxytocin neurons is directly coupled to changes in blood flow velocity and capillary permeability. We further demonstrate that these physiologically elicited vascular changes are regulated by oxytocin, suggesting that oxytocin facilitates its own brain-to-periphery effects.

## RESULTS

### Physiological osmotic challenge increases local neurohypophyseal blood flow

To study neuro-vascular coupling in the HNS, we have devised a live-imaging platform using the zebrafish model. We have previously established that the development, cell biology, and anatomy of the HNS neurovascular interface are conserved between zebrafish and mammals (*13–15*). The core components of larval zebrafish HNS neurovascular interface comprised of elaborate oxytocinergic axonal varicosities, the hypophyseal artery (HyA), which conveys blood flow into a capillary loop (HyC) and a venous hypophyseal outlet (HyV) (**Fig. 1A**). This relatively simple and discernible anatomical architecture allowed us to reproducibly monitor vascular dynamics in the same locale within multiple unanesthetized zebrafish larvae. To monitor blood flow, we employed the *Tg(l-fabp:DBP-EGFP)* reporter (*16, 17*) (**Fig. 1B,C**), in which a serum albumin-like molecule is labeled, allowing visualization and quantification of flow velocities calculated from displacement time kymographs of red blood cells (RBC) (**Fig. S1B**). Capillary diameter was monitored using an endothelial-specific transgenic *Tg(kdrl:EGFP)* reporter (**Fig. S1C**).

**Figure 1.**
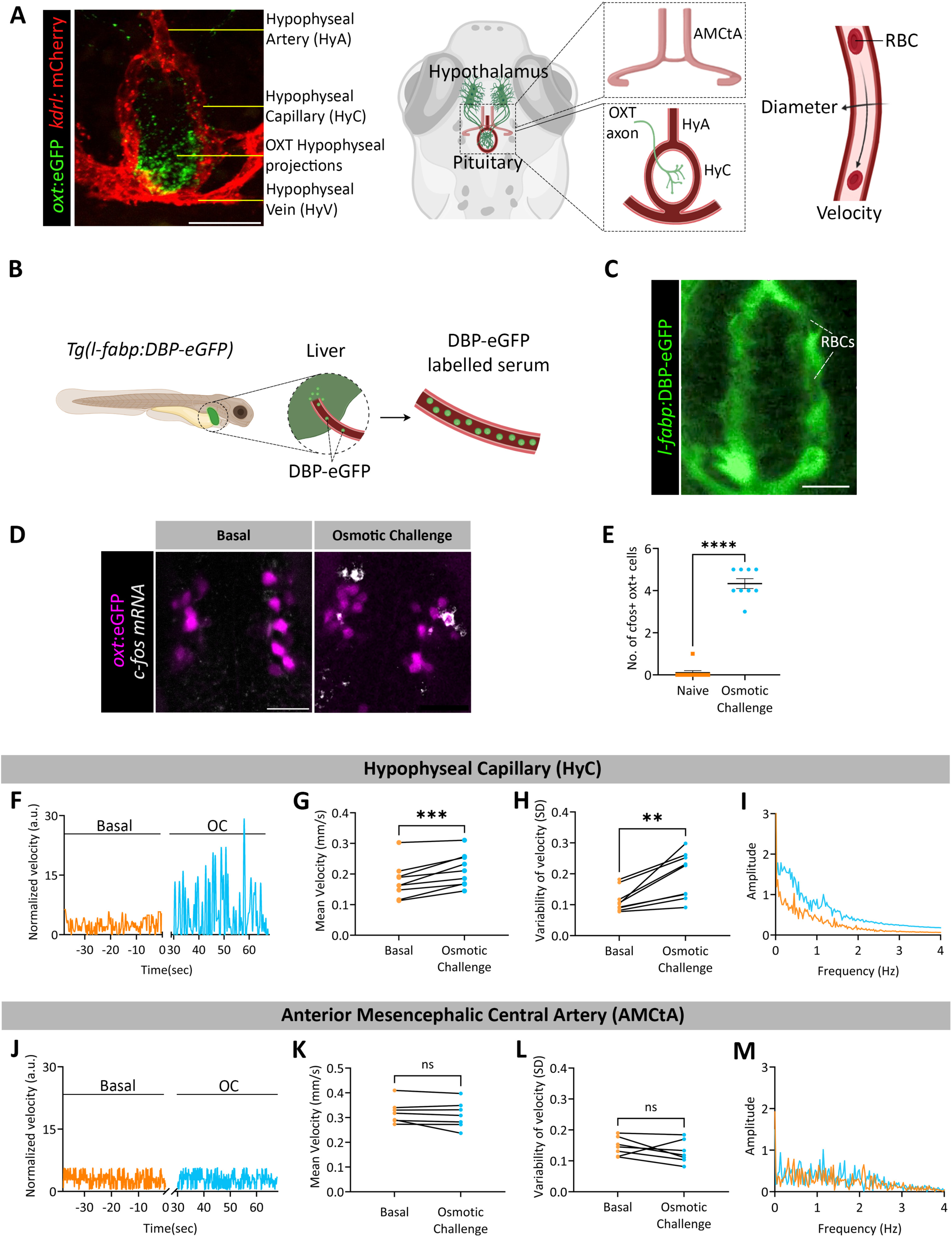
Physiological osmotic challenge increases local hypophyseal blood flow (A) Representative image and scheme showing the hypophyseal neurovascular interface of the transgenic 6-day-old zebrafish larval *Tg(oxt:eGFP;kdrl:mCherry)*. Oxytocin axonal termini and blood vessels are labelled in green and red respectively. (**A, right**) Schema depicting the stereotypic anatomical structure of the hypophyseal artery (HyA), capillary (HyC) and vein (HyV), as well as the anterior mesenchephalic central artery (AMCtA), which was used as a control. Arrows indicate the positions of line scans imaging through and across the vessel were employed to acquire changes in capillary diameter and red blood cell (RBC) velocity traces. (B) Scheme showing the production of DBP-eGFP under the liver promoter, L-fabp, in the *Tg(l-fabp:DBP-eGFP)* transgenic line labelling the serum. (**C**) Snap shot of the hypophyseal capillary loop of the *Tg(l-fabp:DBP-eGFP)* transgenic larvae at 6dpf. Dark shadows are created by the red blood cells (RBCs) on a background of DBP-eGFP labelled serum. (**D-E**) Transgenic *Tg(oxt:eGFP)* zebrafish larvae with labeled oxytocin (OXT) neurons were incubated with hyperosmotic solution equivalent to 50% seawater (17.5 ppt) followed by in-situ hybridization of the activity-dependent *c-fos mRNA.* Representative image of a single z-plane (**D**) showing increased *c-fos* expression in OXT neurons following the osmotic challenge quantified (**E**) as the number of *c-fos* positive OXT neurons expressing cells (p <0.0001, Mann-Whitney test, naïve n=9, OC=10). Scale bar, 20um. (**F**) Representative blood flow traces in the HyC at basal conditions and following the osmotic challenge (OC) which led to increased mean RBC velocities in the HyC (**G,** p=0.001, Paired t-test, n=9) and higher variability of velocities measured by spread of individual velocities around the mean (**H,** p=0.0039, Wilcoxon test, n=9). Single-sided amplitude spectra of the velocities (**I**) indicate an increase in amplitudes for frequencies less than 2Hz following the OC. (**J-M**) Osmotic challenge induced no changes to the blood flow traces (**J**), mean velocity (**K**, p=0.2296, Paired t-test, n=7), variability in velocities (**L**, p=0.2669, Paired t-test, n=7), or any changes in the spectra (**M**) in the AMCtA which served as control for the specific effect of the OC.

To examine whether physiological activation of the HNS altered hypophyseal blood flow, we employed a hyperosmotic challenge known to stimulate OXT and AVP neurons (*18, 19*) (**Fig. S1A**). When zebrafish larvae were acutely transferred from freshwater to hyperosmotic solution equivalent to 50% seawater (17.5 ppt), we observed significant activation of OXT neurons, displaying increased *c-fos* expression in their soma (**Fig. 1D,E**). This hyperosmotic challenge induced a robust increase in blood flow (**Fig. 1F,G**) and variability of RBCs velocities (**Fig. 1H**). The single-sided amplitude spectra, as computed by a Fast Fourier transformation (FFT), indicated that the osmotic challenge induced an increase in the amplitude of velocity fluctuations below 2Hz (**Fig. 1I**).

To ensure that the effect of osmotic challenge on blood flow was specific to the hypophyseal capillary, we measured blood flow in the anterior mesencephalic central artery (AMCtA), which is a well-discernable vascular brain structure located at a similar depth and of similar diameter as the hypophyseal capillary loop (**Fig. 1A**). We found that osmotic challenge did not affect the measured flow parameters of AMCtA vessel (**Fig. 1J-M**). These results show that a physiological challenge, which activates OXT neurons elicits a specific local increase in hypophyseal blood flow velocity.

### The relationship between flow velocity and diameter is determined by HNS vascular geometry

According to Poiseuille’s law for viscous fluids, it was expected that local vasoconstriction of a single capillary would increase flow resistance, resulting in decreased blood flow velocity (*20*). However, we observed that the osmotic challenge, which caused an increase in velocity was accompanied by a decrease in hypophyseal capillary diameter (**Fig. 2A-C**). This effect was specific as the diameter of the adjacent AMCtA capillary did not change following the osmotic challenge (**Fig. 2D-F**). Since it was shown that the geometry of the microvasculature network in the brain plays a crucial role in determining blood flow behaviour (*21, 22*), we resolved the relationship between blood flow velocity and capillary diameter considering the structure of the zebrafish hypophyseal vascular microcircuit. This structure is comprised of a stereotypic arterial inlet, capillary loop, and venous outlet; each having its corresponding length (L), and radius (r) (**Fig. 2G**). We expressed the capillary velocity (V_2_) in terms of the vascular components’ radii and lengths, according to Poiseuille’s law, considering a range of relevant inlet-outlet total pressure differences (Δp) (**Fig. 2G and S1E,** for details, see ‘*Methods*’ section). Plotting the resolved V_2_ values as a function of capillary radii revealed that flow velocity is inversely proportional to an increase in capillary radii that are larger than 3 microns (**Fig. 2H**). This inverse proportionality between capillary velocity and diameter, aligned with our measured values of the hypophyseal capillary radii (3-6 μm **Fig. 2B**). Thus, in accordance with our velocity and radii measurements, we found that for basal velocity to increase after osmotic challenge within a range of relevant estimated inlet-outlet pressure differences (Δp), there must be either a small decrease in radii with a small increase in Δp or a large decrease in radii with no change in Δp (**Fig. 2H’**).

**Figure 2.**
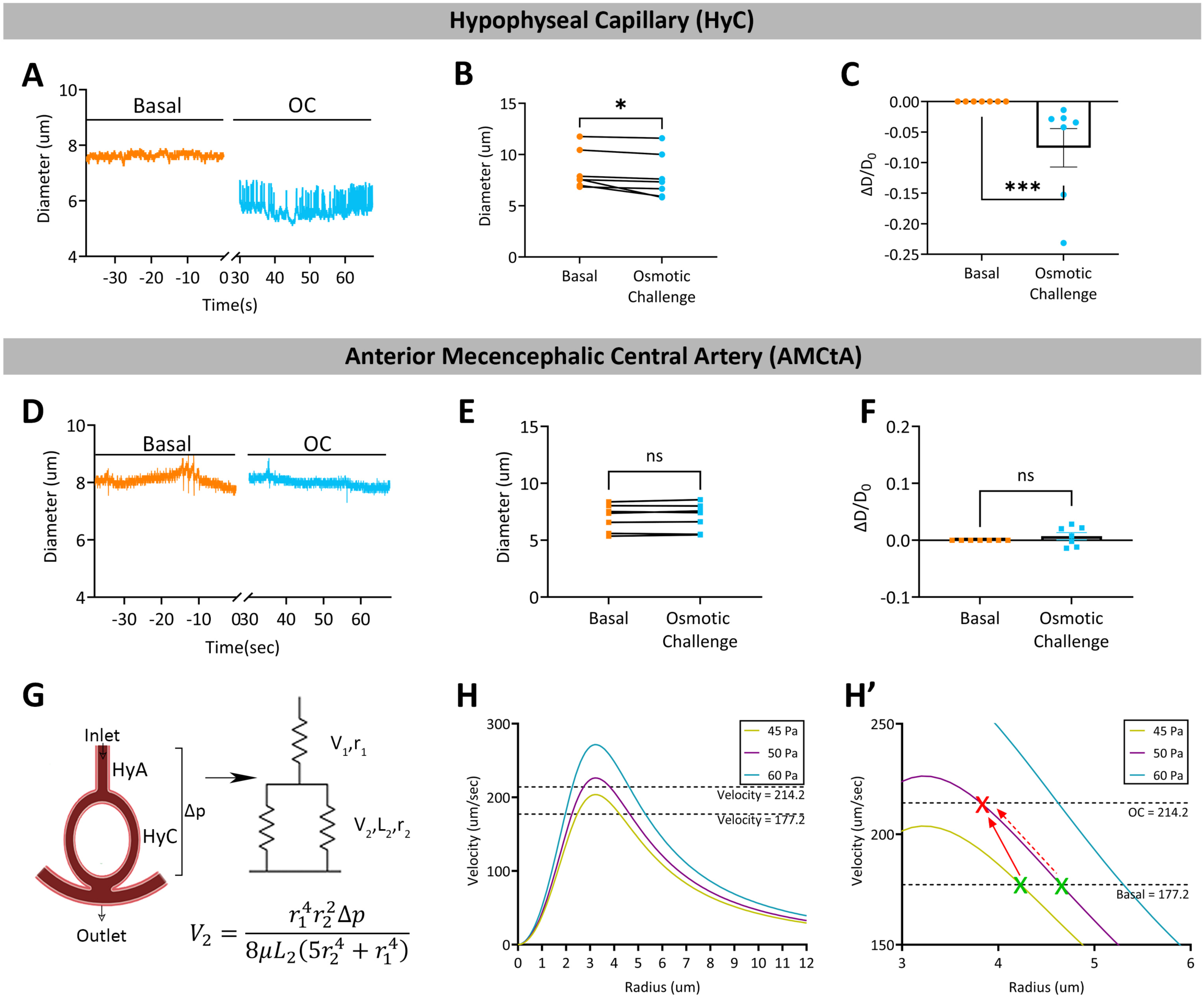
Hypophyseal vascular response is determined by it’s geometry. Changes in capillary diameter in the HyC (**A-C**) and AMCtA (**D-F**). (**A-F**) Representative diameter traces of HyC (**A**) and AMCtA (**D**). Mean diameter of the capillary significantly decreases after OC (**B**, p=0.0156, Wilcoxon test, n=7), as further shown as a ratio of diameter change to basal diameter (**C**, p=0.0006, Mann-Whitney test, n=7). (**D-F**) OC did not affect the mean diameter (**E**, p=0.3076, Paired t-test, n=7), or the ratio of diameter change to basal diameter (**F**, p=0.6900, Mann-Whitney test, n=7) of the AMCtA. (**G**) The predicted relationships between hypophyseal diameter and velocity was theoretically resolved according to Poiseuille’s law for viscous fluids by representing the individual components the hypophyseal vascular microcircuit having an inlet-outlet pressure differences (1′p). The corresponding resistance (R), length (L), and radius (r) of the hypophyseal artery (HyA) and capillary (HyC) are indicated. The resulted equation of hypophyseal flow velocity (V_2_) as a function of the above parameters is shown (Fig. 2G, for details, see ‘*Methods*’ section). (**H**) A range of pressure differences, Δp, was determined by multiple random matches between velocity and radii values that were obtained experimentally. This range was plugged into the resolved V_2_ equation resulting three curves indicating the relationship between velocity and radii of the hypophyseal capillary for a given Δp. The measured average velocities before and after osmotic challenge (OC) are indicated by dashed lines (**H**). The point of intersection between the corresponding basal and OC velocities and the curves for a given Δp are indicated by the green and red Xs (**H’**). A corresponding increase in velocity following an OC requires a small increase in pressure and vasoconstriction (solid line) or a large decrease in radius (**H’**, dashed line).

These results show that an increase in blood flow velocity can be caused by vasoconstriction given the geometry of the hypophyseal vascular microcircuit.

### Hypophyseal blood flow is regulated by oxytocin signaling

We tested whether the observed neuronal and vascular responses to an osmotic challenge are coupled. This was demonstrated using the *Tg(oxt:Gal4;UAS:NTR-mCherry)* genetic system for conditional cell ablation (**Fig. 3A**). Specific death of OXT cells expressing the nitroreductase (NTR) enzyme was induced by the administration of the drug Nifurpirinol (NFP), which is converted to a cytotoxic agent by NTR (*23–25*). Treatment of these transgenic animals for 18 hours with NFP led to a significant decrease in OXT-positive hypophyseal axonal projections (**Fig. 3B,C**). When subjected to an osmotic challenge, control Gal4/UAS-negative larvae displayed increased blood flow velocity while their sibling larvae with ablated OXT neurons did not (**Fig. 3D,E**). The mean capillary blood flow velocity, amplitudes, and frequency spectra, and variability in velocity did not show a significant difference after the osmotic challenge of OXT ablated larvae (**Fig. 3F-I**). Notably, whereas the osmotic challenge caused vasoconstriction in non-ablated controls, it led to vasodilation of hypophyseal capillaries in larvae with ablated OXT neurons, presumably as a response to the absence of OXT neurovascular innervation and low basal OXT secretion (**Fig. 3J-M**). The effects of this genetic OXT-ergic ablation were confined to the hypophyseal vasculature as they were not observed in the control AMCtA vessel (**Fig. 3N-P**).

**Figure 3.**
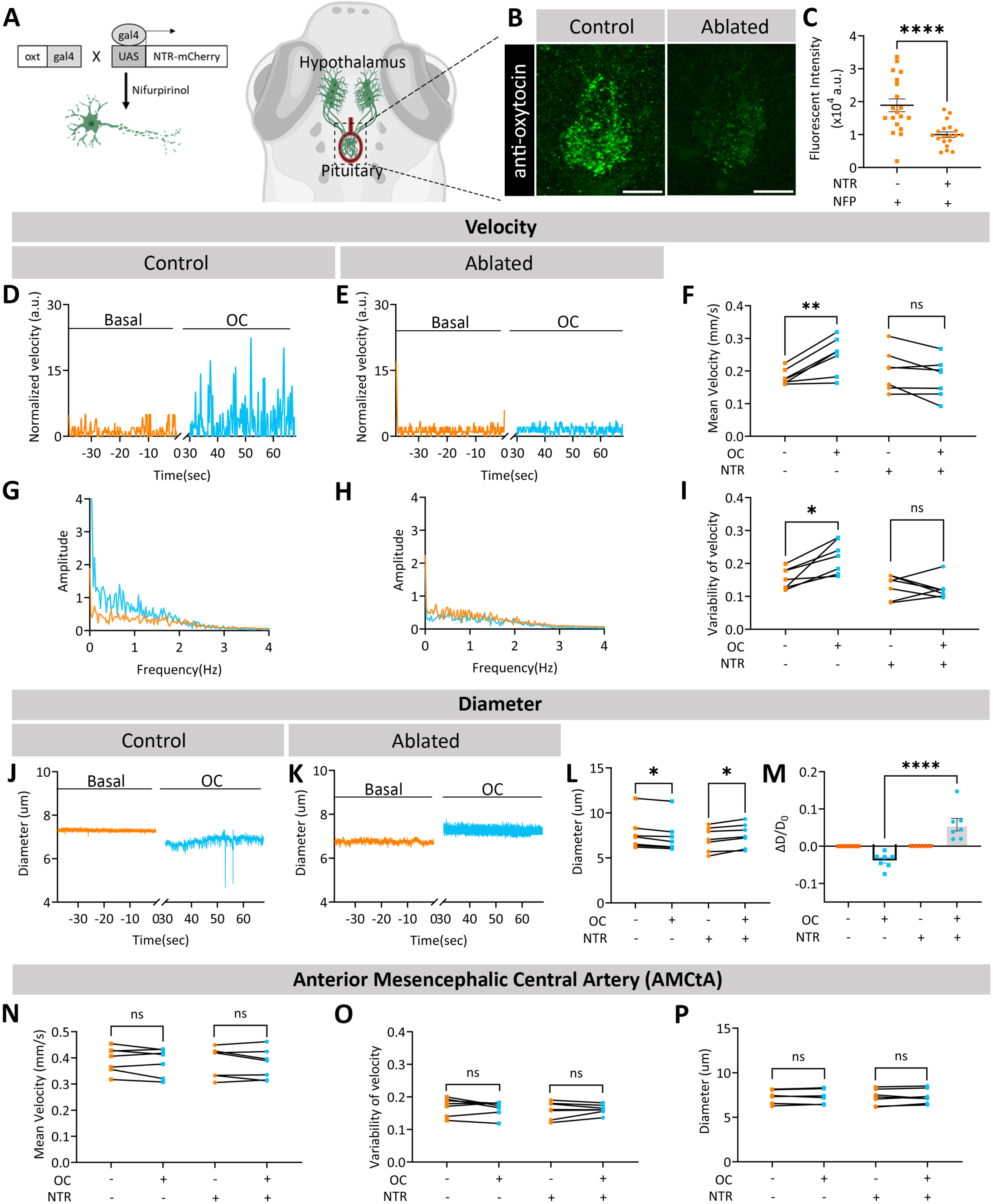
Ablation of oxytocin neurons attenuates vascular response to an osmotic challenge. (**A**) Schema representing the transgenic *Tg(oxt:gal4;UAS:NTR-mCherry)* larvae, in which oxytocin neurons express the nitroreductase (NTR) enzyme which reduces the drug nifurpirinol (NFP) into a cytotoxic compound causing specific oxytocinergic cell death. (**B-C**) *Tg(oxt:gal4;UAS:NTR-mCherry)* larvae were treated for 18 hours with 5uM NFP and immune-stained with and antibody directed to endogenous OXT protein. NTR-(control) and NTR+ (Ablated) siblings were treated with NFP, followed by anti-OXT antibody staining. Representative confocal z-stacks images show a reduction of OXT protein in neurohypophyseal axonal projections following NFP treatment of NTR-positive larvae (**B**). Scale bar, 10um. Quantification of OXT-immunoreactive fluorescence shows significantly reduction of OXT in neurohypophyseal termini in the NTR+ cohort (**C**, p<0.0001, Mann-Whitney test, control n=19, treated n=19). Blood flow velocity traces representing the osmotic challenge (OC) increase in hypophyseal blood flow velocity in control (**D**), but not OXT-ablated (**E**) larvae. (**F**) Mean velocity after OC was increased in NTR-larvae (p=0.0046, Paired t-test, n=7), but not in their NTR+ siblings (p=0.0947, Paired t-test, n=7). (**G, H**) Single sided amplitude spectra of the NTR+ cohort show no difference before and after OC as compared to the controls. Variability in velocity was increased after OC in NTR-cohort also (**I**, p=0.0081, Paired t-test, n=7), but not in their NTR+ siblings (**I**, p=0.5781, Wilcoxon test, n=7). (**J-K**) Representative diameter traces before and after osmotic challenge showing capillary vasoconstriction of control NTR-larvae (**J**) compared to vasodilation of ablated NTR+ siblings (**K**). **(L)** Following OC, the hypophyseal capillary of NTR-larvae showed a decrease in diameter (p= 0.0156, Wilcoxon test, n=7) while NTR+ larvae showed an increase in diameter (p=0.0045, Paired t-test, n=7). The ratio between diameter change to basal diameter after OC (**M**) was significantly different between the NTR- and NTR+ cohorts (p<0.0001, Dunn’s multiple comparison, NTR-n=7, NTR+ n=7). (**N-P**) Mean velocity (p=0.3941, Paired t-test, n=7), variability of velocity (p=0.6983, Paired t-test, n=7) and diameter (p=0.5249, Paired t-test, n=7) of the AMCtA did not change after OC in NTR+ larvae treated with NFP. The control NTR-larvae treated with NFP also did not show any difference in mean velocity (p=0.3095, Paired t-test, n=7), variability of velocity (p=0.5781, Wilcoxon test, n=7) and diameter (p=0.4430, Paired t-test, n=7) in the AMCtA after OC.

In view of our observed effects of OXT neuron ablation, we hypothesized that OXT itself might regulate local hypophyseal blood flow velocity upon its release. We found that endothelial cells isolated from zebrafish pituitaries express mRNAs encoding the two known functional OXT receptors, *oxtra,* and *oxtrb,* with markedly higher expression levels of *oxtrb* (**Fig. S2**). We therefore examined whether the administration of a selective oxytocin receptor antagonist, L-368,899, which is known to inhibit both zebrafish OXT receptors (*26, 27*), affects hypophyseal blood flow. We found that larvae treated with L-368, 899, were refractory to the osmotic challenge in terms of challenge-induced changes in mean flow velocity parameters (**Fig. 4A-F**) and capillary diameter (**Fig. 4G-J**). Administration of L-368, 899 did not influence blood flow velocity in the extra-hypophyseal AMCtA blood vessel ruling out a general systemic effect of the antagonist (**Fig. 4K-M**).

**Figure 4.**
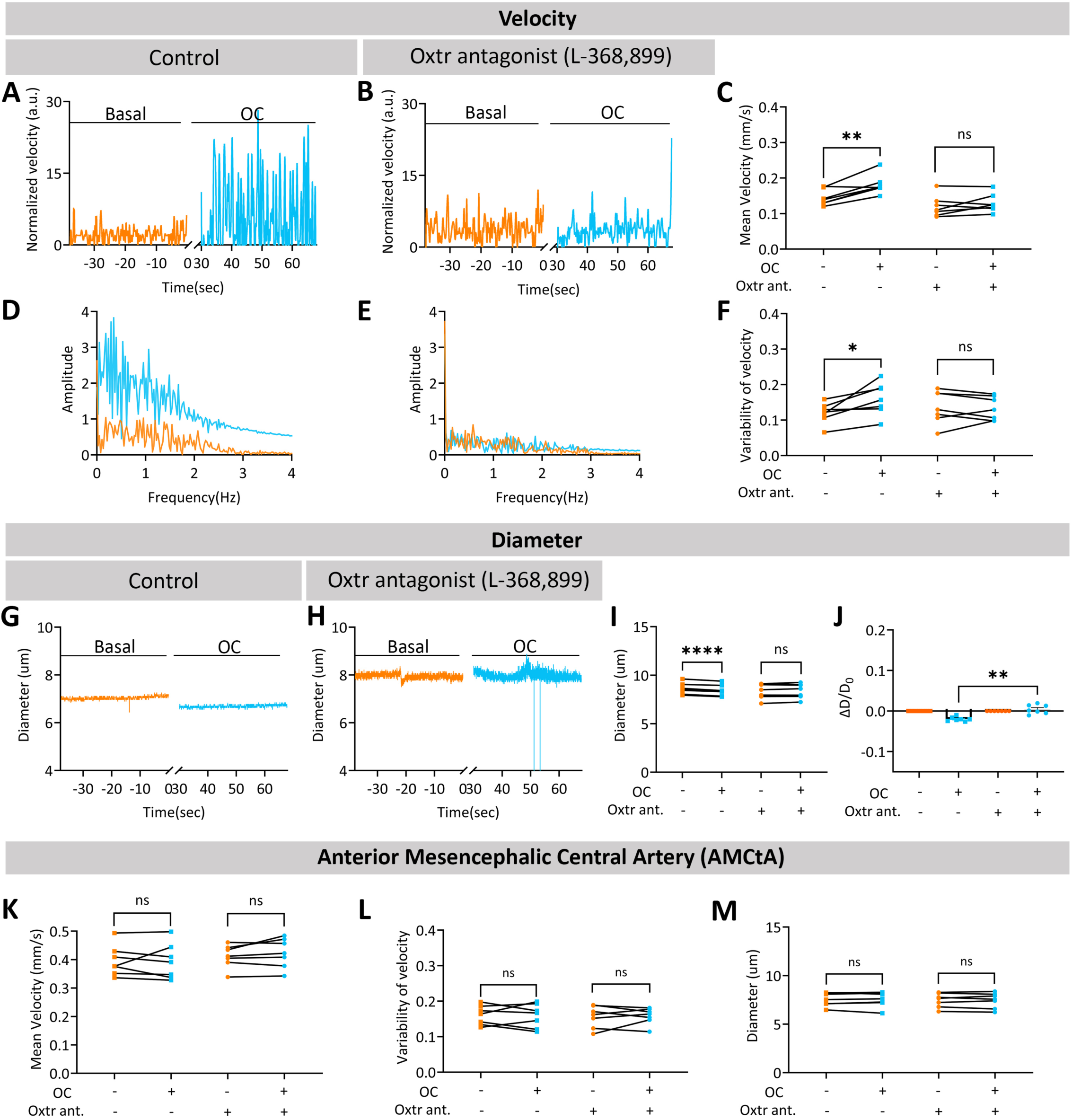
Hypophyseal blood flow is regulated by oxytocin signaling (**A, B**) Representative hypophyseal blood flow velocity traces before and after osmotic challenge (OC). Larvae treated with the Oxtr antagonist (Oxtr ant.) L-368,899 show no change in response to OC as compared to their control sibling. Mean velocity (**C**) after OC in the oxtr ant-cohort show an increase (p=0.0033, Paired t-test, n=7), that is not observed in L-368,899 antagonist treated siblings (p=0.2509, Paired t-test, n=7). (**D, E**) Single sided amplitude spectra show that while the frequency spectra of a control cohort were affected the OC, larvae treated with the Oxtr antagonist show no difference. Variability in velocities (**F**) was increased following OC (p=0.0213, Paired t-test, n=7), but not in Oxtr antagonist-treated siblings (p=0.7195, Paired t-test test, n=7). (**G, H**) Representative diameter traces showing vasoconstriction following osmotic challenge of control sibling but not in larvae treated with Oxtr antagonist (**I**) No significant change in the mean capillary diameter of the Oxtr antagonist-treated larvae which were subjected to OC (p=0.3892, Paired t-test, n=7) compared to a decrease in diameter in the control cohort (p=0.0001, Paired t-test, n=7). (**J**) The ratio between diameter change to basal diameter after OC showed a significant difference between the control and antagonist-treated cohorts (p=0.0010, Dunn’s multiple comparison, Oxtr ant-n=7, Oxtr ant+ n=7). (**K-M**) Mean velocity (p=0.2008, Paired t-test, n=7), variability of velocity (p=0.8040, Paired t-test, n=7) and diameter (p=0.8509, Paired t-test, n=7) of the AMCtA did not change after OC in Oxtr ant+ larvae. The control Oxtr ant-larvae also did not show any difference in mean velocity (p=0.8573, Paired t-test, n=7), variability of velocity (p=0.8533, Paired t-test, n=7) and diameter (p=0.9422, Paired t-test, n=7) in the AMCtA after OC.

Next, we tested whether direct activation of OXT neurons, rather than through osmotic challenge, is sufficient to modulate hypophyseal blood flow and diameter. To stimulate OXT neurons, we employed the CoChR-tdTomato channelrhodopsin variant, an efficient optogenetic tool for zebrafish (*28*). Injection of *UAS:CoChR-tdTomato* construct into the OXT specific driver line, *Tg(oxt:Gal4)* (*29*), resulted in specific expression of the CoChR-tdTomato protein in OXT neurons (**Fig. 5A,B**). Exposure of *Tg(oxt:Gal4;UAS:CoChR-tdTomato)* larvae to blue light resulted in a transient but significant increase in blood flow velocity, which peaked at approximately 30-50 seconds following light exposure (**Fig. 5D, E**). The control CoChR-tdTomato-negative siblings did not respond to blue light (**Fig. 5C, E**). Light-activated CoChR-positive larvae exhibited a marked increase in 1′V/V_0_ as compared to their CoChR-negative siblings (**Fig. 5F**). The responses of hypophyseal vasculature to optogenetic stimulation of OXT neurons were dependent on oxytocin receptor signalling as the above effects were significantly attenuated by the administration of the OXT receptor antagonist, L-368, 899 (**Fig. 5G-I**). In accordance with the vasoconstrictive response of the hypophyseal capillary to osmotic challenge, light activation of CoChR-positive, but not CoChR-negative, OXT neurons caused a reduction in mean capillary diameter across time (**Fig. 5J-M**). As in the case of the increased flow velocity, CoChR-induced vasoconstrictive response was inhibited by L-368, 899 (**Fig. 5N-P**). Notably, hypophyseal vascular responses (i.e., flow and diameter) to optogenetic activation of OXT neurons were transient as compared to the hyperosmotic challenge. Although this may imply the existence of additional mediators of the effects of osmotic challenge on hypophyseal vascular dynamics, it could be due to the artificial activation of OXT neurons through optogenetics, as opposed to a genuine physiological stimulus, which activates these neurons.

**Figure 5.**
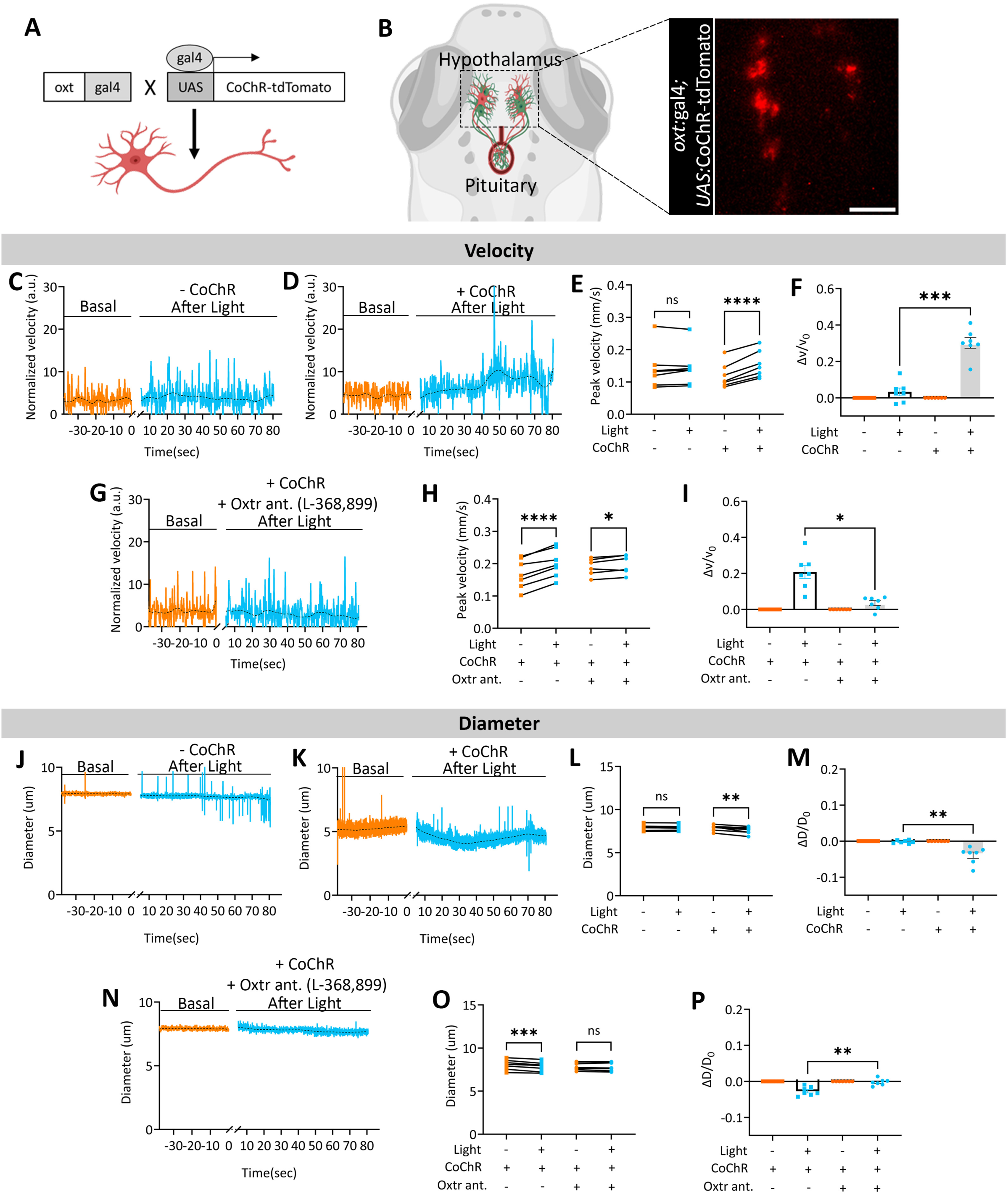
Optogenetic stimulation of oxytocin neurons increases hypophyseal blood flow and is dependent on oxytocin signaling (**A-B**) Scheme representing the Gal4-UAS system for expression of a blue light-activated channel rhodopsin (*UAS:CoChR-tdTomato*) in oxytocin (OXT) neurons using *Tg*(oxt:Gal) transgenic driver. (**B**) Representative single z-plane confocal image of oxytocin neurons expressing CoChR-tdTomato. Scale bar, 20um. (**C, D**) Representative blood flow velocity traces before and after light stimulation in larvae expressing CoChR show an increase in blood flow velocity as compared to their control sibling. (E) Blue light stimulation of CoChR+ larvae led to an increase in mean peak blood flow velocity (p=<0.0001, Paired t-test, n=7). Control CoChR-siblings were not responsive to the light stimulation (p=0.4688, Wilcoxon test, n=7). (F) The ratio between velocity change to basal velocity after light stimulation of the CoChR+ cohort increased as compared to their CoChR-sibling cohort (p=0.0225, Dunn’s multiple comparison, CoChR-n=7, CoChR+ n=7). (G) Representative blood flow velocity traces before and after light stimulation in larvae expressing CoChR and treated with the Oxtr antagonist L-368,899 (Oxtr ant.). (H) Mean velocity after light stimulation in the CoChR+ Oxtr ant+ larvae show an increase (p=0.0313, Wilcoxon test, n=7), which was also observed in their CoChR+ Oxtr ant-siblings (p=<0.0001, Paired t-test, n=7). (**I**) The ratio between velocity change to basal velocity after light stimulation in the vehicle-treated CoChR+ was significantly larger as compared to their CoChR+ siblings treated with L-368,899 (p=0.0287, Dunn’s multiple comparison, Oxtr ant-n=7, Oxtr ant+ n=7). (**J-K**) Representative diameter traces before and after light stimulation in larvae expressing CoChR show an increase in blood flow velocity as compared to their control sibling. (**L-M**) Mean diameter after light stimulation in the CoChR+ larvae show a decrease (p=0.0043, Paired t-test, n=7), that is not observed in their CoChR-siblings (p=0.3349, Paired t-test, n=7). The ratio between diameter change to basal diameter after light stimulation in the CoChR+ cohort is lower as compared to their CoChR-sibling cohort (p=0.0047, Dunn’s multiple comparison, CoChR-n=7, CoChR+ n=7). (N) Representative diameter trace before and after light stimulation in larvae expressing CoChR and treated with the Oxtr antagonist L-368,899. (O) Light stimulation of CoChR+ larvae led to a decrease in mean capillary diameter (p=0.0010, Paired t-test, n=7). The capillary diameter of the antagonist-treated CoChR+ larvae did not respond to the optogenetic activation (p=0.6043, Paired t-test, n=7). (P) The ratio between diameter change to basal diameter after light stimulation of the antagonist-treated CoChR+ larvae were significantly lower as compared to vehicle-treated CoChR+ sibling cohort (p=0.0060, Dunn’s multiple comparison, Oxtr ant-n=7, Oxtr ant+ n=7). Dashed lines in all representative traces indicate a LOWESS curve fitted to the data with a smoothing window of ten points.

These results show that local hypophyseal vascular responses triggered by a hyperosmotic physiological challenge are coupled to OXT neuronal activation and are mediated by OXT receptor(s).

### Hypophyseal vascular permeability is regulated by OXT signaling

We hypothesized that OXT-mediated neurovascular-coupling may affect the vascular passage of neurohormones and other proteins by regulating diffusion rate through the fenestrated hypophyseal capillaries (*16*). We visualized and quantified hypophyseal capillary permeability using the *Tg(l-fabp:DBP-EGFP)* reporter described above (**Fig. 6A**). We performed fluorescent recovery after photo-bleaching (FRAP) followed by line scan imaging comparing the fluorescent recovery of bleached and unbleached regions of interest positioned on either side of the hypophyseal capillary (**Fig.6A’, B**). Paired comparisons of permeability before and after the challenge indicated that the average time taken to recover half the initial fluorescent intensity (T_1/2_) following an osmotic challenge was significantly shorter than in the basal condition, indicating a higher rate of diffusion through the hypophyseal capillary (**Fig. 6C**).

**Figure 6.**
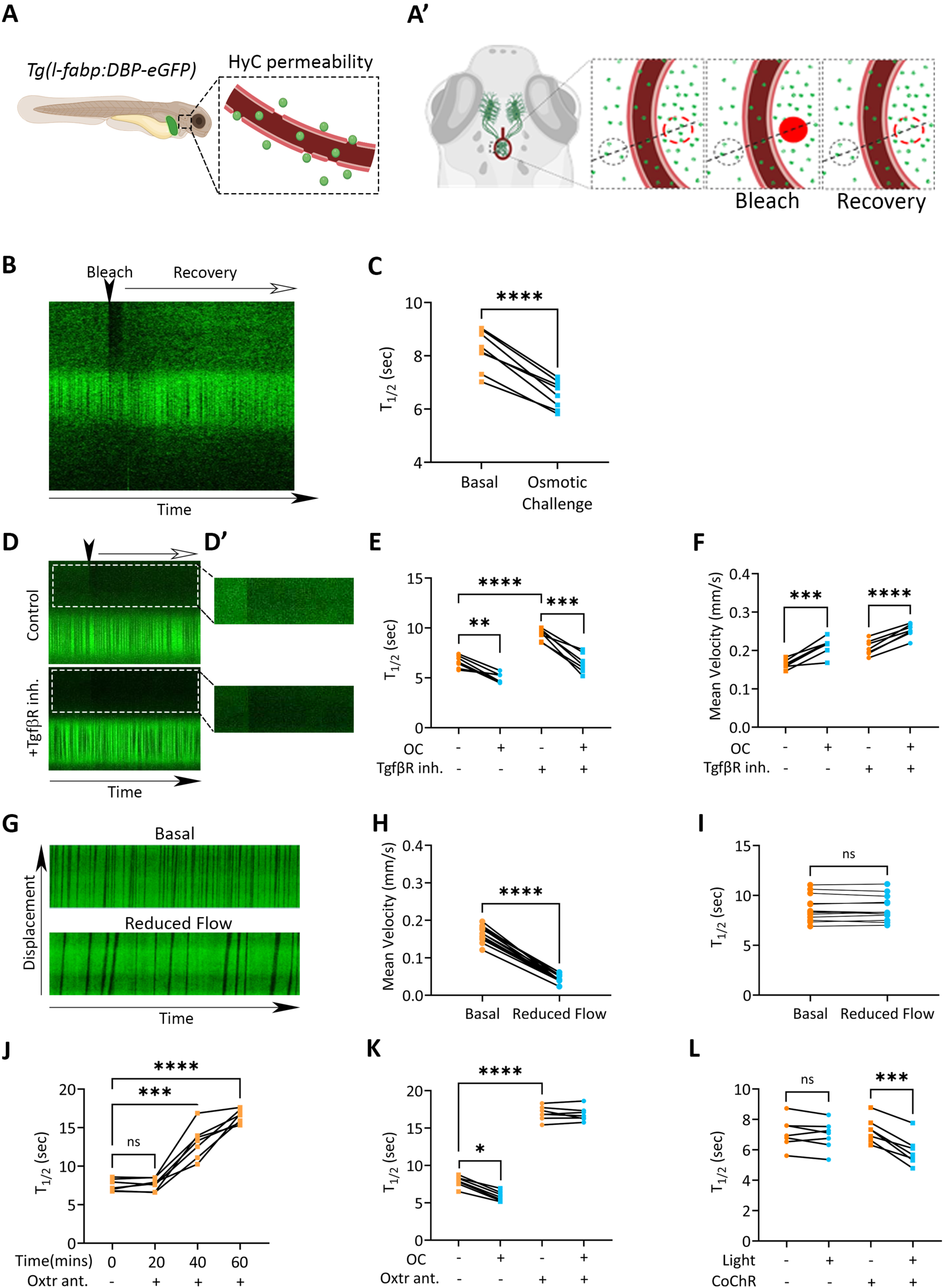
Hypophyseal vascular permeability is regulated by oxytocin signaling (**A**) Graphical representation of the permeable hypophyseal capillary visualized in the *Tg(l-fabp:DBP-eGFP)* larvae. (**A’**) Scheme depicting the hypophyseal capillary permeability assay in which fluorescent recovery after photobleaching (FRAP) of the extravasated DBP-EGFP signal the *Tg(l-fabp:DBP-EGFP)* reporter was measured in a region of interest (ROI) within the hypophyseal capillary loop (red circle) compared to a control ROI outside of the loop. (**B**) A kymograph representing a line scan across time showing the bleached and unbleached ROIs on either side of the hypophyseal capillary, which is marked by dashed lines. The bleaching time point is marked by an arrowhead. (**C**) Average FRAP recovery half-time T_1/2_ was decreased after osmotic challenge (OC) (p<0.0001, Paired t-test, n=8). (D) Kymographs representing FRAP traces of naïve (unchallenged) larva displaying vehicle-treated control larva and larva treated with 25 μM of the Tgfβ receptor inhibitor SB431542 (TgfβR in.). The bleaching time point is marked by an arrowhead. Higher intensity images of the FRAP kymographs before and after photobleaching are shown on the right (**D’)**. (E) The basal T_1/2_ values of SB431542-treated larvae were significantly higher than their vehicle-treated siblings (p<0.0001, Dunnett’s multiple comparison, TgfβR inh.-n=7, TgfβR inh.+ n=7). Larvae treated with SB431542 exhibit a decrease in T_1/2_ values after OC (p=0.0002, Paired t-test, n=7) as do their vehicle-treated control larvae (p=0.0012, Paired t-test, n=7). (F) Mean blood flow velocity values following OC was increased in larvae that were treated with the TgfβR inhibitor (p<0.0001, Paired t-test, n=7) as do their vehicle-treated controls (p=0.0007, Paired t-test, n=7). (G) Representative kymographs of larvae exhibiting basal versus reduced blood flow. (H) Application of 1 mg/ml of α-bungarotoxin for 30 seconds immobilizes larvae without affecting blood flow velocity (basal), whereas incubation with α-bungarotoxin for 150 seconds reduced mean blood flow velocity values (p<0.0001, Paired t-test, n=13). (I) Reduction in blood flow velocity did not affect capillary permeability as indicated by FRAP measurements of T_1/2_ values (p=0.0854, Paired t-test, n=13). (J) Hypophyseal capillary permeability was monitored at 20 mins intervals before and after the application of Oxtr antagonist. A significant increase in T_1/2_ after 40 and 60 mins of antagonist treatment (p=0.8090, Basal *vs*. 20min; p=0.0004, Basal vs 40min; p<0.0001; Basal *vs*. 60min, Dunnett’s multiple comparison, n=7). (K) The increment in capillary permeability which is induced by OC (p<0.0001, Paired t-test, n=7) is blocked in larvae which were treated with the Oxtr antagonist L-368,899, showing no change in T_1/2_ after OC (p=0.6147, Paired t-test, n=7). In accord with Fig. 6J above, basal T_1/2_ values of L-368,899-treated larvae were significantly higher than their untreated siblings (p<0.0001, Dunnett’s multiple comparison, Oxtr ant-n=7, Oxtr ant+ n=7). (L) Optogenetic activation of OXT neurons was performed using oxt:Gal;*UAS:CoChR-tdTomato* transgenic larvae, as described in the legend to Figure 5 following by FRAP-based permeability measurements. Blue light stimulation of CoChR+ larvae led to a decrease in T_1/2_ values (p=0.0003, Paired t-test, n=7). T_1/2_ values were unaffected by light in CoChR-larvae (p=0.2559, Paired t-test, n=7).

We then investigated the inter-dependent relationship between blood flow velocity and permeability. We have previously shown that Tgf-β signaling regulates hypophyseal vascular permeability (*13*). We now show that albeit the observed attenuation of basal hypophyseal permeability by the selective TGF-β receptor inhibitor, SB431542 (**Fig. 6D, E**), blocking Tgf-β signaling did not affect either the basal blood flow velocity or the response to osmotic challenge (**Fig. 6F**). We next examined whether decreasing blood flow affects permeability. We observed that treatment of zebrafish larvae with the cholinergic receptor inhibitor alpha-bungarotoxin (α-BTX) for 150 seconds led to a nearly 50% reduction in blood flow velocity (**Fig. 6G-H**). This reduction in blood flow had no effect on hypophyseal vessel permeability (**Fig. 6I)**. These results indicate that hypophyseal vascular permeability and blood flow velocity are mutually independent.

To determine the involvement of OXT signaling in hypophyseal vascular permeability, we performed the FRAP analysis in the presence of the OXT receptor antagonist. Basal permeability was measured continuously at different time points following the administration of the OXT receptor antagonist. A significant increase in T_1/2_ was observed up to an hour after the administration of the antagonist, indicating a cumulative decrease in protein diffusion rate through the hypophyseal capillary (**Fig. 6J**). Furthermore, the permeability-enhancing effect of OC was also attenuated in antagonist-treated larvae as compared to their untreated siblings (**Fig. 6K**).

The involvement of OXT in modulating permeability was further determined by direct optogenetic activation of OXT neurons combined with FRAP analysis. A significant decrease in T_1/2_ was measured in light-activated, CoChR-tdTomato-expressing larvae compared to their inactivated basal state (**Fig. 6L**). The control CoChR-negative larvae showed no response to light (**Fig. 6L**). Thus, direct activation of oxytocin neurons enhances hypophyseal permeability.

Taken together, our data show that hyperosmotic physiological challenge elicits an increase in local blood flow velocity and permeability of the hypophyseal vasculature, where OXT is released into the peripheral circulation. These hypophyseal vascular responses are influenced by the local vascular geometry and are coupled to OXT neuronal activation, which is mediated by OXT receptors. We submit that OXT may function as a self-perpetuating hormone that facilitates its own peripheral uptake by regulating blood flow dynamics.

## DISCUSSION

Neurohormones are key factors interfacing between the brain and periphery, yet not much is known about their behavior at the site of release, and in particular, the factors affecting their uptake into the blood. We focus here on the evolutionarily conserved HNS, which is a canonical neuroendocrine interface wherein neurohormones are directly released from the brain into the blood. The data we have gathered indicates that when faced with a hyperosmotic physiological challenge, there’s a noticeable increase in hypophyseal blood flow velocity and vascular permeability where OXT is released into the peripheral circulation. The local hypophyseal vascular responses to OXT neuronal activation are influenced by vascular geometry and mediated by OXT receptors. This leads us to propose that OXT may function as a self-perpetuating hormone, regulating its own peripheral uptake by influencing blood flow dynamics and permeability.

### Neurovascular coupling in the HNS

Upon stimulation of OXT neurons, OXT is released from the nerve termini into the hypophyseal perivascular space and thereafter exerts effects on peripheral target tissues. However, there is a gap in our understanding of the exact events that bring about efficient transport of OXT to the periphery. Unlike the rapid, short-distanced post-synaptic actions of neurotransmitters in the brain, OXT must travel a long distance through the dense hypophyseal perivascular space and its concentration in the blood must increase rapidly in response to physiological demands. This suggests that additional factors may be required for an efficient coordination between the release of neurohormones and their peripheral effects. We hypothesized that activity-mediated axonal release needs to be coupled to blood flow to regulate an effective transfer of OXT into the systemic circulation. However, despite extensive understanding of the electrical activity of oxytocin (OXT) neurons, the role of local blood flow in coordinating neurohormone secretion remains unclear (*30, 31*). Lafont et al. inferred the existence of such a mechanism, demonstrating that the injection of growth hormone-releasing hormone (GHRH) leads to increased local blood flow, coinciding with the timing of the growth hormone (GH) pulse in the peripheral circulation (*32*). These authors discussed the possibility that flow dynamics of the pituitary microvasculature network may influence the pulsatile secretion of growth hormone (*32*).

We show here that an osmotic challenge stimulates OXT neuronal activity concomitant with a specific local increase in volumetric blood flow in the hypophyseal capillary abutting OXT axons termini. This result is in agreement with the reported increased neurohypophyseal blood flow in chronically water-deprived rats (*33, 34*). Others have recently shown a connection between osmotic challenge and changes in supraoptic nucleus (SON) blood flow (*35*). They reported that an acute systemic salt load activates AVP neurons, decreasing blood flow and causing temporary hypoxia, which intensifies the excitatory response of AVP neurons to the systemic hyperosmotic salt challenge (*35*).

Local regulation of organ perfusion rate relies on local changes in vascular resistance, which is controlled by capillary radius (*36, 37*). Thus, local vasoconstriction of a single capillary, obeying Poiseuille’s flow, will increase resistance and decrease flow (*20*). Having said that, the geometry of a particular microvasculature network, including factors such as capillary length, branching patterns and positions of inlet-outlet, plays an important role in determining flow velocity (*22, 38*). In the case of the specific configuration of the hypophyseal vascular circuit, we found that blood flow velocity increases upon vasoconstriction. This result also fits our theoretically resolved relationship between blood flow velocity and capillary diameter, considering the specific architecture, lengths, and radii of the hypophyseal vascular components.

### Role of oxytocin signaling

Vascular endothelial cells were shown to express the OXT receptor and have been implicated in the regulation of vascular tone and blood pressure in mammals and fish (*39–43*). It was shown that OXT can exert both vasodilation and vasoconstriction via differential activation of either endothelial nitric oxide or ERK signaling (*39*). We show here that, as in other species, zebrafish OXT neurons are osmoresponsive, and the effect of osmotic challenge on hypophyseal blood flow is mediated by OXT signaling. Similarly, increased osmolality, which was induced by dehydration, led to an increased blood flow in the rat neurohypophysis (*33, 34*). Using Brattleboro rats, Kapitola et al. concluded that the increase in hypophyseal blood flow following dehydration could not be mediated by AVP as these rats cannot synthesize this neurohormone (*34*). However, it is likely that OXT is accountable for the observed dehydration effect on Brattleboro rat hypophysis. This is supported by our findings that the impact of hyperosmotic challenge is mediated by OXT signaling, and that direct optogenetic stimulation of OXT neurons influences both blood flow and diameter.

We show that endothelial cells express OXT receptors and that blocking OXT signaling attenuates the induced changes in blood flow and capillary diameter. However, it is important to note that other cell types in the neurohypophysis may also play a role in OXT-mediated vascular response. For example, pericytes have been implicated in the regulation of blood flow in the brain (*44–46*). Additionally, hypophyseal astroglial pituicytes, which receive synaptoid connections from HNS axon termini, facilitate the release of OXT and AVP to the periphery in response to parturition, lactation, and dehydration (*29, 47, 48*).

### Hypophyseal vascular permeability

Although the HNS is considered part of the central nervous system, its vascular component has unique properties that differ from neurovascular units in other brain areas. Unlike most BBB-containing vessels of the CNS, neurohypophyseal capillaries are fenestrated, allowing bi-directional transfer of HNS hormones and blood-borne substances between the brain and circulation (*13, 16, 49*). Our research shows that protein diffusion through the hypophyseal capillary increases after a hyperosmotic challenge. Although we show that the increase in hypophyseal blood flow velocity and permeability appears to be independent of one other, both occur in an oxytocin-dependent manner. Thus, blocking OXT signaling decreases hypophyseal permeability and direct activation of OXT neurons enhances permeability.

Modulation of local blood flow has been linked to hormonal uptake in several instances. For example, somatostatin causes vasoconstriction, reducing blood flow and increasing vascular permeability of pial vessels of the parietal cortex (*50*). Blood flow in the median eminence, which contains fenestrated capillaries, influences food intake by regulating the access of ghrelin to the arcuate nucleus (*51, 52*). Finally, pericytes, which regulate blood flow (*45, 46*) were shown to regulate vascular permeability to circulating leptin (*53*). We propose that in the case of the neurohypophysis, oxytocin orchestrates an efficient neuroendocrine secretion machinery by affecting both hypophyseal blood flow velocity and permeability, thereby coupling its own release and transport into the blood.

### Stimulus-secretion-uptake coupling

Based on the results we have presented, we propose the following paradigm, which we term ‘*stimulus-secretion-uptake coupling*’: When HNS neurons are stimulated, OXT is released into the hypophyseal parenchyma and establishes a concentration gradient of OXT between the parenchyma and the local capillary. The diffusion of OXT into the capillary tends to bring the system closer to equilibrium, thus reducing the diffusion of oxytocin into the vessel. The increase in local volumetric flow, which is induced by OXT, has a dual function: **1**) enhancing local transport of OXT to the periphery and **2**) increasing diffusion of the secreted OXT into the blood by maintaining a local oxytocin concentration gradient. This, together with the increased permeability, which is also evoked by OXT, ensures its efficient transfer into the peripheral circulation in response to physiological demands.

## ACKNOWLEDGEMENTS

We thank Takashi Kawashima and Pablo Blinder for valuable discussions and advice; Roy Hofi, Estar Regev and fish facility personnel. G.L. lab is supported by the Israel Science Foundation (#349/21); Hedda, Alberto, and David Milman Baron Center for Research on the Development of Neural Networks; Wolfe Family Center for Research on Neuroimmunology and Neuromodulation; Sagol Center for Research on the Aging Brain; and Maurice and Vivienne Wohl Biology Endowment and Foundation for Higher Education and Culture. P.R. was supported by a research grant for a student fellowship from the Benoziyo Endowment Fund for the Advancement of Science and by the Weizmann–CNRS Collaboration Program. G.L. is an incumbent of the Elias Sourasky Professorial Chair.

## AUTHOR CONTRIBUTIONS

Conceptualization, P.R. and G.L.; Methodology, P.R. Investigation, P.R., O.R. and G.L.; Resources, P.R., Formal analysis, P.R. and O.R.; Writing-original draft, P.R. and G.L.; Writing-review and editing, P.R., G.L., and O.R.; Visualization, P.R.; Funding acquisition, G.L.; Project administration, P.R. and G.L.

## MATERIALS AND METHODS

### Husbandry and housing conditions of experimental animals

All experiments using zebrafish were approved by the Weizmann Institute’s Institutional Animal Care and Use Committee (Application number 00590123-3 and 16970919-3). Zebrafish were maintained and bred using standard protocols and according to FELASA guidelines.

### Transgenic animals

The following transgenes were used in this study: *Tg(l-fabp:DBP-eGFP)*,*Tg(kdrl:eGFP), Tg(oxt:gal4-VP16; UAS:NTR-mCherry; l-fabp:DBP-eGFP/kdrl:eGFP), Tg(oxt:gal4-VP16; UAS:CoChR-tdTomato; l-fabp:DBP-eGFP/kdrl:eGFP).* Larvae that were to be imaged were kept in the Danieau’s buffer with 0.003% PTU from 24hpf till the day of the experiment.

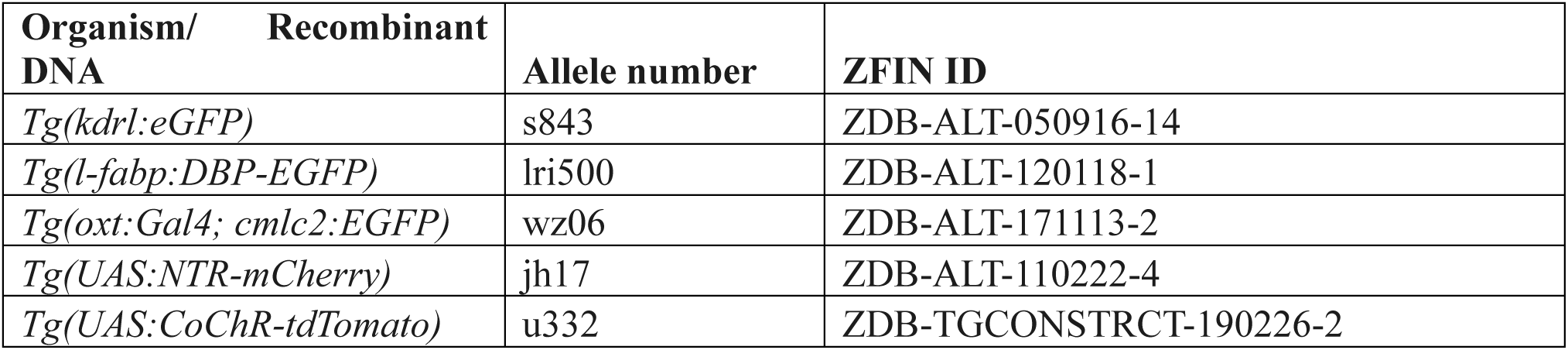

### *In-vivo* imaging of live zebrafish

Six days post fertilization (dpf) zebrafish larvae were immobilized with a 30s incubation with 1mg/ml alpha-bungarotoxin followed by a quick wash in E3 media and mounted in 0.8% low-melt agarose in a custom moulded 3cm plate filled with 2% agarose. The plate was filled with 3ml E3 media prior to imaging. Samples were excited using the Zeiss LSM7 Multi-Photon Microscope with a pulsed laser (Coherent Chameleon) tuned to 870nm set at a 4-6% laser intensity (max. 210mW) through a 20x water immersion Pln Apochromatic Lens. Emitted light was detected through the BiG detector after passage through a BP500-550 filter. Line scans were obtained by manually defining a single-pixel scan path (pixel scaling of 0.03um) as required which was then scanned for a pre-defined number of frames using the Galvanometric scanning head.

### Osmotic challenge, cell ablation and pharmacological treatments

Zebrafish were osmotically challenged using 50% instant ocean salt (17.5 gr/l; Instant Ocean Sea Salt, Aquarium Systems) dissolved in E3 media (5.03mM NaCl, 0.17mM KCl, 0.33mM CaCl_2_.2H_2_O, 0.33mM MgSO_4_.7H_2_O). Oxytocin neurons were ablated by treating 5 dpf *Tg(oxt:gal4-VP16; UAS:NTR-mCherry)* larvae on the background of either *Tg(l-fabp:eGFP)* or *Tg(kdrl:eGFP)* with 5uM Nifurpirinol (NFP; LGC, 13411-16-0) for 18h at 28.5^0^C in E3 media. Larvae where then washed for two hours in E3 media before live imaging. Controls for the experiment were *Tg(oxt:gal4-VP16; UAS:NTR-mCherry)* negative siblings which were also treated with NFP to account for the effects of the drug itself.

Oxytocin receptor signaling was attenuated by treating 6 dpf *Tg(l-fabp:eGFP)* or *Tg(kdrl:eGFP)* larvae with 100mM Oxtr antagonist L-368,899 (No.2641, Tocris Bioscience) dissolved in E3 media for 1hour at 28.5^0^C followed by live imaging.

Tgfβ receptor inhibition was carried out by treating 4 dpf *Tg(l-fabp:eGFP)* or *Tg(kdrl:eGFP)* larvae with 25uM SB431542 (S4317, Sigma-Aldrich) dissolved in DMSO for 48h at 28.5^0^C followed by live imaging.

To slow down blood flow in the brain/pituitary, 6 dpf *Tg(l-fabp:eGFP)* or *Tg(kdrl:eGFP)* larvae were incubated in a-bungarotoxin for 150 seconds followed by a wash and live imaging.

### Optogenetic activation of oxytocin neurons

Adult Tg(oxt:gal4-VP16) were crossed with Tg(l-fabp:eGFP) or Tg(kdrl:eGFP) and the resultant embryos were injected with 40ng/ul of UAS:CoChR-tdTomato (Addgene; plasmid #124233) construct and 40ng/ml of Transposase mRNA at the single-cell stage. Larvae were visually sorted at 5d pf for strong expression of tdTomato in the oxytocin neurons under a fluorescent stereo microscope. Control larvae were siblings that were not injected with the UAS:CoChR-tdTomato construct. All experimental cohorts were grown in an incubator at 28.5^0^C in the dark.

Hypophyseal capillary was imaged under basal conditions, with minimal light activation of the neurons, followed by 10 minutes of rest. Oxytocin neurons were then stimulated for 5 seconds with a 470nm Fiber-coupled LED (ThorLabs Inc,USA) through a fiber optic cannula (400um, 0.39NA; ThorLabs) placed near the head of the larvae at 1.5 mW/mm2, followed immediately by live imaging of the hypophyseal capillary only.

### Monitoring and analysis of blood flow velocity

Arbitrary line scans with a length of approx. 20 um along the vessel length were obtained at 133 Hz in the *Tg(l-fabp:DBP-eGFP)* transgenic line. These line scans were then combined into a multi-frame time-displacement image (kymograph). Two signals were derived from two different points on the displacement axis, and along the time axis. These signals were a representation of the fluctuation in fluorescence intensities across time. Independently, each of these signals gives a measure of the number of red blood cells (RBCs) flowing through the cross-sectional area of the hypophyseal capillary per unit time. On comparing the two signals from each RBC, taken at points x1 and x2, we get the displacement of the RBC within the time frame between t1 and t2. Using these coordinates, the slope of each RBC was computed, and converted to velocity (**Fig. S1C)**. The individual RBC velocities in the given kymograph were then plotted as a function of normalized displacement. With an optimal sampling rate of RBCs per second, individual velocities of the RBCs across time were used to obtain a power density spectrum to identify the frequencies at which the flow velocities oscillate. Changes between basal and treated conditions were also depicted as a change in power of particular frequencies or a shift in the frequency itself using a fast Fourier transform (FFT).

### Capillary diameter measurements

To measure capillary diameter, line scans were taken perpendicular to vessel length at 133 Hz in the *Tg(kdrl:eGFP)* line. The length of the line was approx. 40um, which covered both ends of arms of the hypophyseal capillary. These line scans were then combined into a multi-frame time-displacement image (kymograph), where each line perpendicular to the x-axis represents a single diameter trace at the corresponding time (**Fig. S1D)**. Signals derived from each line corresponding to time values on the x-axis were plotted to obtain two Gaussians each representing the left or right vessel wall. Inner and outer diameter measurements were obtained by measuring the distance between the inner and outer Full-Width Half-Maxima (FWHM) values of these Gaussians. This diameter measured is then converted from pixels to microns and plotted as the final graph as the diameter changes across time for both basal and treated conditions.

### Vascular permeability assay

Permeability was visualized and quantified using the *Tg(l-fabp:DBP-EGFP)* reporter by employing fluorescence recovery after photobleaching (FRAP) to determine the average recovery half-time (T_1/2_) of the extravasated DBP-EGFP signal. This was carried out by defining two circular regions of interest (ROIs), each of 5μm in diameter, on either side of the hypophyseal capillary. A line scan path (pixel size 0.06um) was drawn perpendicular to direction of flow, through the vessel, and through the centers of the two ROIs. Basal measurements of raw fluorescence were obtained, followed by bleaching the ROI in the area within the hypophyseal loop (Fig. 6A). Photobleaching was performed at 100% laser power (approx. 3500mW) for 2.2s at 870nm, followed by the acquisition of the recovery of fluorescence along the line scan path. Raw fluorescence was extracted directly by Zen and was exported to MATLAB to calculate T_1/2_ values. T_1/2_ was calculated by fitting an exponential curve to the recovery florescence and calculating the intercept on the time axis at the point of 50% of the maximal fluorescence. The unbleached ROI was used as a control for unintentional bleaching, and basal fluorescence was used to validate the maximal fluorescence values after recovery.

### Whole-mount immuno-fluorescence, HCR mRNA in-situ hybridization

Six day-old larvae were fixed in 4% PFA, followed by PBST washes and overnight incubation with 100% Methanol at −20C. Larvae were washed with PBST (0.5% Tween-20 in 1x Phosphate Buffered Saline), incubated in blocking solution for 1 hour at room temperature, followed by overnight incubation at 4C with Guinea Pig anti-oxytocin antibody (Peninsula Laboratories International Inc., CA, USA, no. T-5021) in a blocking solution (5% goat serum in PBST). Larvae were washed with PBST, and then incubated for 4 hours at room temperature with Mouse anti-guinea pig Cy5 secondary antibody (Jackson ImmunoReasearch Laboratories, PA, USA) followed by PBST washes.

Hybridization chain reaction (HCR) in-situ v3.0 probes for *c-fos mRNA* were custom designed and synthesized by Molecular Instruments (USA). In-situ hybridization was performed according to Chen et al. (*15*).

Larvae were then mounted in 75% glycerol and imaged with a 40x oil immersion lens using the Zeiss LSM800 confocal setup using the appropriate emission/excitation filters.

### FACS Sorting and qRT-PCR

Five adult pituitaries of the *Tg(kdrl:eGFP)* transgenic line were dissected into Ca^2+^ free Ringer’s solution (116 mM NaCl, 2.9 mM KCl, 5.0 mM HEPES, pH 7.2), followed by a 30 minute incubation in 1X TrypLE Express (Gibco, 12604-013). Pituitaries were then mashed onto a 40um cell stainer and flushed into an Eppendorf using 500ul suspension buffer (L15 media without phenol red (Gibco, 21083-027), 1% FBS, 0.5% pen-strep solution). 25ul of 20ug/ml DAPI was added, and the cells were sorted into three populations: all pituitary cells (GFP+ and GFP-), endothelial cells only (GFP+), and endothelial depleted (GFP-). populations. Cells were collected in a lysis buffer (0.3% NP-40, 0.1% FBS, 1% RNase Inhibitor) and converted into cDNA directly (Takara, #RR036A). qRT-PCR was performed with 200cells per well and three technical triplicates using the following gene-specific primers:

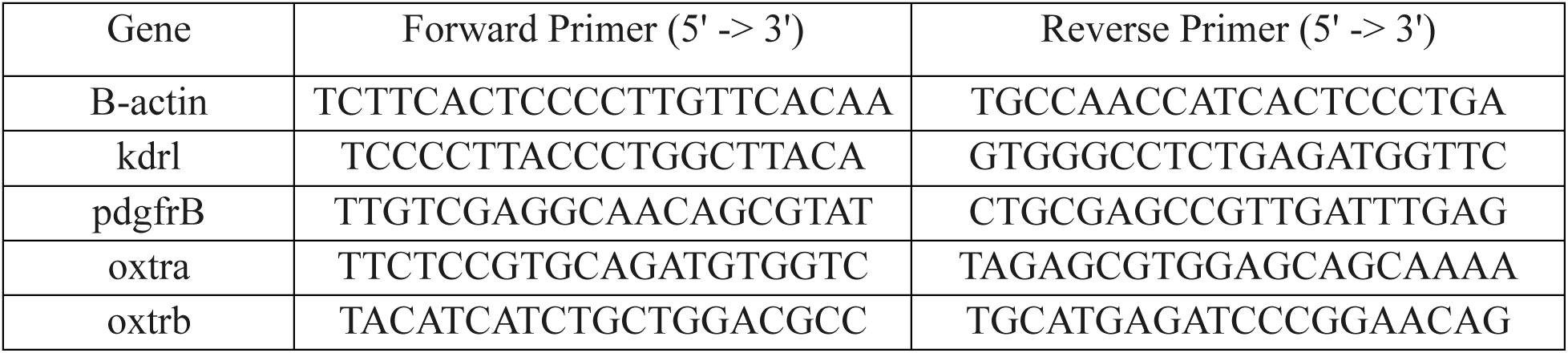

### Codes

All algorithms to measure velocity, diameter and permeability were custom written on MATLAB (Mathworks, USA) and are available on request.

### Statistical analysis

All statistical analyses and graphing were performed using PRISM 10.2 (GraphPad Software Inc, San Diego, USA). Data tables were exported from MATLAB to Prism were required. The normality of the data sets were tested using the Shapiro-Wilk test, and parametric/non-parametric tests were carried out with either a Paired t-test, Mann-Whitney test or the Wilcoxon test for paired samples. All the tests were two-tailed analyses. Welch’s ANOVA with Geisser-Greenhouse corrections or Kruskal-Wallis tests was carried out where appropriate with multiple comparisons (Dunnett’s, Dunn’s or Tukey’s). Significance thresholds for all analyses were set at a 0.05 threshold and p-values were corrected for multiple comparisons were necessary. No data points were excluded as outliers in all analyses.

### Theoretical resolution of hypophyseal vascular response

Expressing Poiseuille’s law for fluid flow through a vessel in terms of flow velocity and vessel radius, we get:

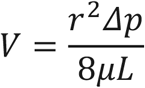

where r is the radius of the vessel, V is the velocity, Δp is the pressure difference between the two ends of the vessel, µ is the viscosity, and L is the length of the vessel (*54*).

To account for the geometry of the vascular microcircuit, we reduced the hypophyseal vasculature as a function of their vascular resistances (Fig. 2G).

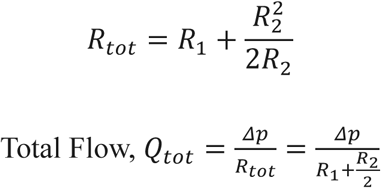

where R_tot_, R_1_, and R_2_ are the total resistance, arterial and capillary resistance respectively, and Q_tot_ is the total flow rate.

Pressure difference in the capillary, Δ*p*_2_ = *Q*_tot_*R*_par_

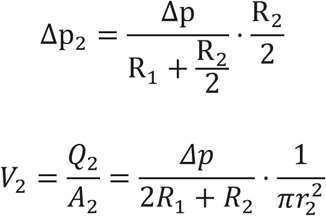

where R_par_ is the total resistance on the parallel arms of the hypophyseal capillary

Breaking down the denominator and expressing resistance in terms of radius, we get:

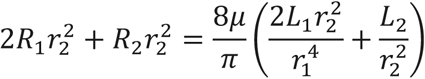

where r_1_and r_2_ are the radii of the artery and capillary, L_1_ and L_2_ are the lengths of the artery and capillary, respectively.

Substituting the above equation back in the original equation, and L_1_ / L_2_ = 2.5L_2_ (experimentally derived):

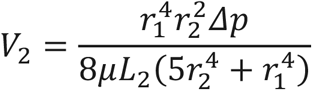

From this equation, given our experimental data, we derived a range of Δp values by pair matching different combinations of mean velocity and radii from out experimental cohorts. A velocity vs radius graph was then plotted using a range of radii with the above pre-defined sets of values for each curve (Fig. 2H).

---

**Supplemental Fig. S1.**
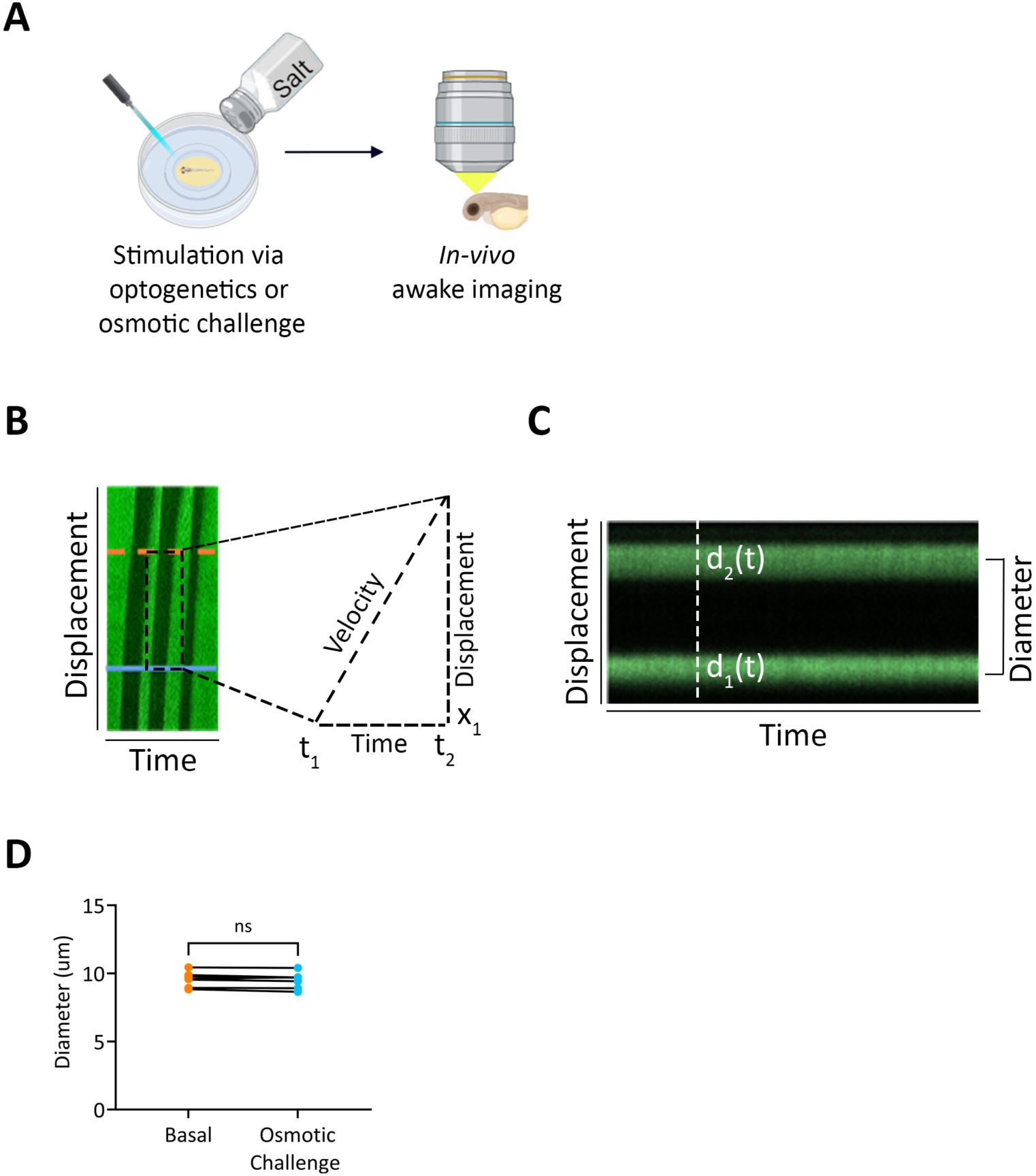
(A) Graphical depiction of experimental workflow where either osmotic challenge or optogenetic stimulation is administered to the zebrafish larvae followed by live *in-vivo* awake imaging. (B) Kymograph shows red blood cells (RBCs) as black streaks on a background of serum labelled with DBP-eGFP. Signals from each RBC, taken at points x1 and x2, denote the displacement of the RBC within the time frame between t1 and t2. The velocity is given by the tangential derived from these components. (**C**) A kymograph across the blood vessel where each line perpendicular to the x-axis represents a single diameter trace at the corresponding time. d_1_(t) and d_2_(t) represent the position of the inner walls of the vessel on the displacement axis for a given time, t. (**D**) Diameter of the hypophyseal artery shows no change after OC (p=0.1783, Paired t-test, n=7).

**Supplemental Fig. S2.**
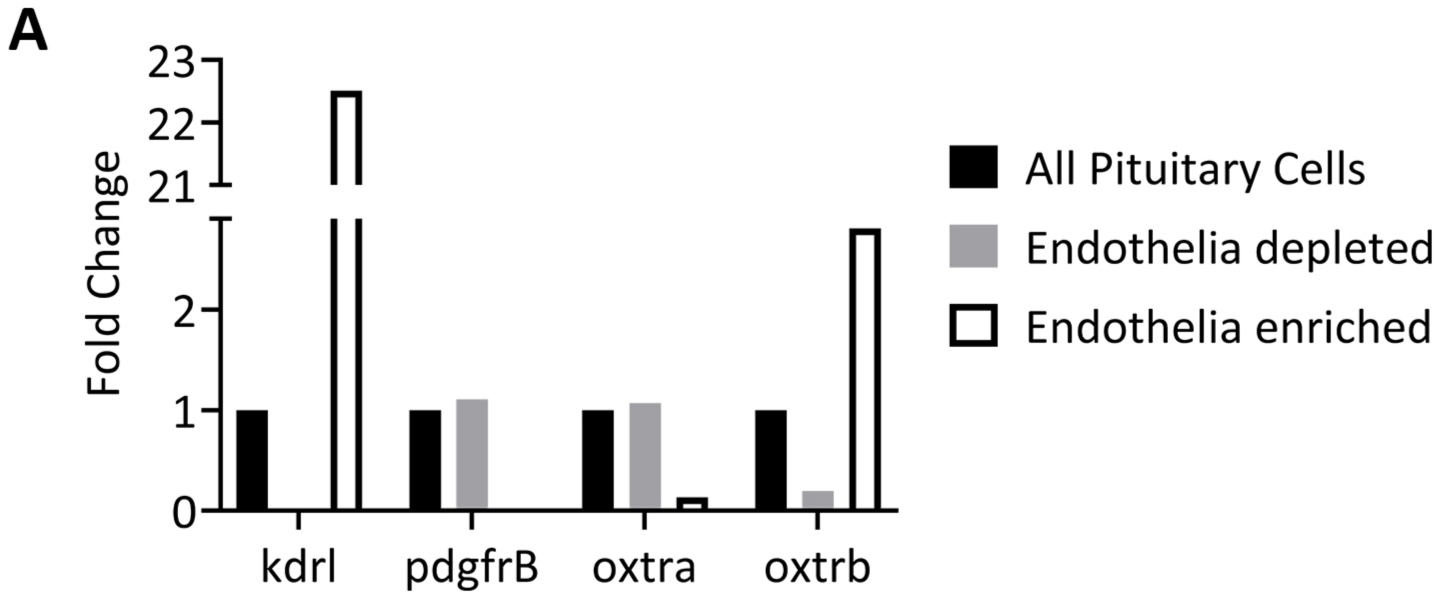
(**A**) Adult pituitary sorted for all cells, endothelia only, and endothelia depleted populations show an enrichment of *oxtrb* mRNA in the endothelial population. *oxtra* mRNA was enriched in the endothelial depleted population but was still observed in low levels in the endothelia population. All comparisons of fold change are made with respect to the mRNA levels of the genes in the entire pituitary.

